# A potent neutralizing nanobody against SARS-CoV-2 with inhaled delivery potential

**DOI:** 10.1101/2020.08.09.242867

**Authors:** Junwei Gai, Linlin Ma, Guanghui Li, Min Zhu, Peng Qiao, Xiaofei Li, Haiwei Zhang, Yanmin Zhang, Yadong Chen, Weiwei Ji, Hao Zhang, Huanhuan Cao, Xionghui Li, Rui Gong, Yakun Wan

**Affiliations:** Shanghai Novamab Biopharmaceuticals Co., Ltd., Shanghai 201318, China; Shanghai Key Laboratory of Molecular Imaging, Shanghai University of Medicine and Health Sciences, Shanghai 201318, China; CAS Key Laboratory of Special Pathogens and Biosafety, Wuhan Institute of Virology, Center for Biosafety Mega-Science, Chinese Academy of Sciences, Wuhan, Hubei 430071, China; Laboratory of Molecular Design and Drug Discovery, School of Science, China Pharmaceutical University, Nanjing 211198, China

**Keywords:** Nanobody, SARS-CoV-2, receptor-binding domain, variants, neutralizing activity, large-scale production, nebulization, inhalation

## Abstract

The outbreak of COVID-19 has emerged as a global pandemic. The unprecedented scale and severity call for rapid development of effective prophylactics or therapeutics. We here reported Nanobody (Nb) phage display libraries derived from four camels immunized with the SARS-CoV-2 spike receptor-binding domain (RBD), from which 381 Nbs were identified to recognize SARS-CoV-2-RBD. Furthermore, seven Nbs were shown to block interaction of human angiotensin converting enzyme 2 (ACE2) with SARS-CoV-2-RBD-variants, bat-SL-CoV-WIV1-RBD and SARS-CoV-1-RBD. Among the seven candidates, Nb11-59 exhibited the highest activity against authentic SARS-CoV-2 with ND_50_ of 0.55 μg/mL. Nb11-59 can be produced on a large-scale in *Pichia pastoris*, with 20 g/L titer and 99.36% purity. It also showed good stability profile, and nebulization did not impact its stability. Overall, Nb11-59 might be a promising prophylactic and therapeutic molecule against COVID-19, especially through inhalation delivery.

**Graphical Abstract:** 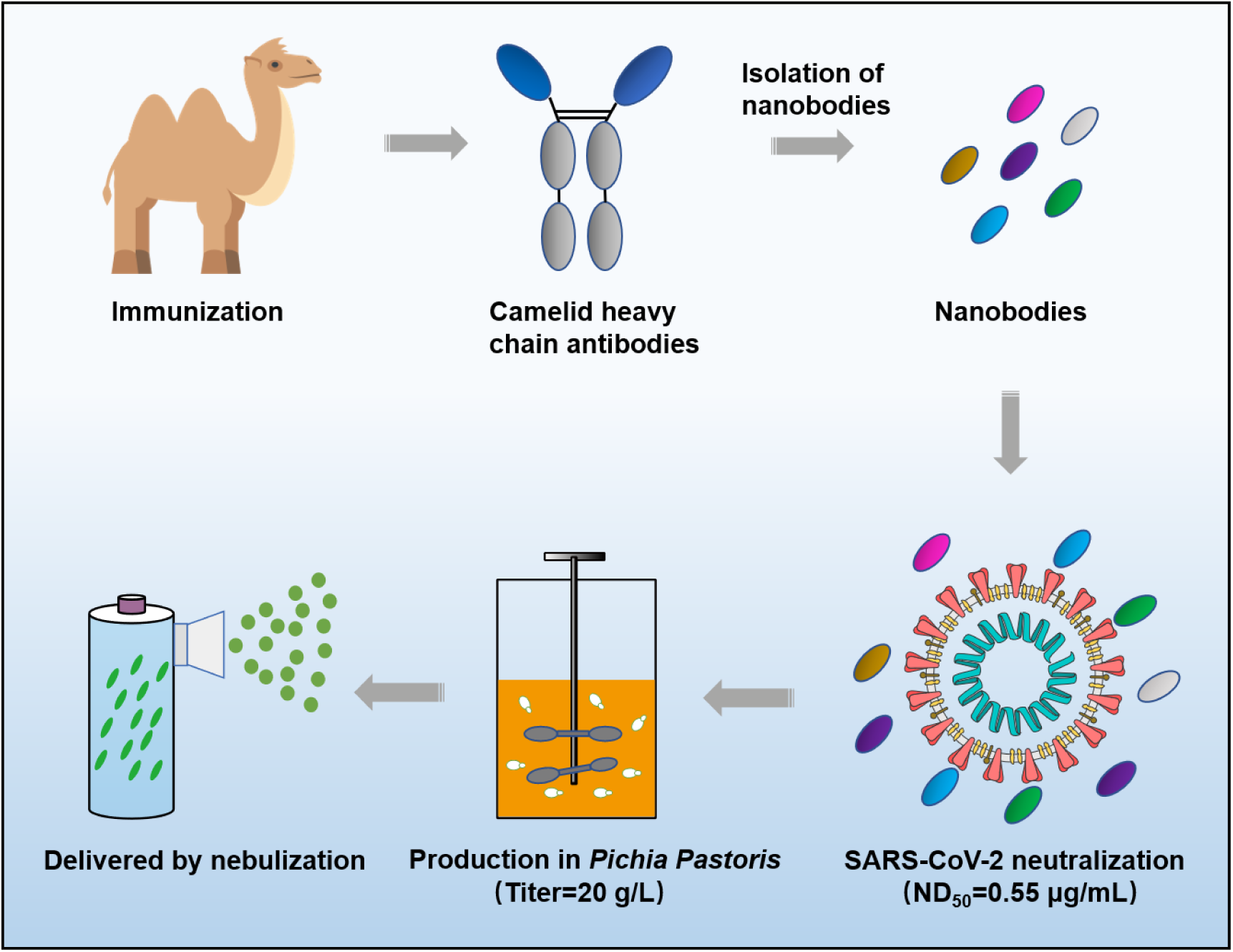

## Introduction

The world-wide spread of coronavirus disease 2019 (COVID-19) caused by SARS-CoV-2 is the third major coronavirus outbreak in the past 20 years. SARS-CoV-2 is closely related to SARS-CoV, several SARS-like bat CoVs and pangolin coronaviruses^1-3^. However, SARS-CoV-2 demonstrates a higher human-to-human transmissibility than SARS-CoV. Patients with confirmed SARS-CoV-2 infection had mild to severe respiratory illness with symptoms of fever, cough, headache, dyspnea and pneumonia^4^. To date, no vaccine or therapeutic has been approved to prevent or treat COVID-19. As of August 1^st^, there are more than 17 million confirmed cases of COVID-19, resulting in more than 680,000 related deaths globally. COVID-19 remains a huge challenge to public health. Therefore, it is urgent to develop prophylactic vaccines or therapeutic drugs to provide solutions for future pandemic.

Similar to other coronaviruses, the spike glycoprotein (S) homotrimer on the SARS-CoV-2 plays a pivotal role in receptor binding and viral entry^5^. The spike glycoprotein is segregated into two functional subunits, termed S1 and S2. The S1 subunit is responsible for the binding of host cell receptor via the interaction between its C-terminal receptor-binding domain (RBD) and human angiotensin converting enzyme 2 (ACE2). S2 subunit plays an important role in fusion of the viral and cellular membranes. The receptor binding, proteolytic processing and structural rearrangement of S protein are regarded as the key events during the merging of the viral envelope with host membranes^6,7^. Crystal structure of SARS-CoV-2-RBD in complex with human ACE2 has been solved, which provides the basis for development of SARS-CoV-2-targeting vaccines and therapeutics^8^.

Currently, a large quantity of drugs and vaccines have been developed or under development for the prevention and treatment of COIVD-19. Biologic drugs neutralizing SARS-CoV-2-RBD/ACE2 interaction, especially, are growing explosively. There are at least 9 SARS-CoV-2-RBD antibodies or combination of antibodies being evaluated in clinical trials, with REGN-COV2 being the most advanced (NCT04452318, Phase 3)^9,10^. It is reported that the resistance mutation in the virus can be induced under the selection pressure of drugs^11^. It is thus an important direction to develop COVID-19-targeting antibodies with broad-spectrum activities against both the wild-type and the mutant viral strains. Besides, novel delivery strategies of antibodies are encouraged, such as antibodies suited for inhalation, which are convenient and can be widely used for COVID-19 prevention.

In addition to conventional monoclonal antibodies (mAbs), heavy-chain-only antibodies (HCAbs) isolated from camels provide an alternative paradigm in development of therapeutic antibodies. HCAbs consist of only two heavy chains without light chains, thereby only containing a single variable domain (VHH) referred to as a single-domain antibody or Nb in the absence of the effector domain^12,13^. In addition to having affinities and specificities for antigens similar to those of traditional antibodies, Nbs show smaller size and higher stability than most antibodies^14,15^. Due to their structure nature, Nbs can be easily constructed into multivalent or multi-specific formats and produced with convenient steps of purification at low manufacturing cost^16,17^. Given these biophysical advantages, Nbs can be easily nebulized and delivered directly to lungs via an inhaler, which make Nbs a particularly promising format for developing neutralizing antibodies targeting respiratory pathogens including SARS-CoV-2^18,19^. The inhaled formulation could potentially allow for easier administration outside the hospital, at earlier stages of disease, which is very important in helping to stem the tide of the pandemic caused by COVID-19.

Here, we report Nb phage display libraries derived from four camels immunized with the SARS-CoV-2 spike RBD, from which 381 Nbs were identified to recognize the RBD including the variants of SARS-CoV-2. Unlike the reported Nbs that require fusion with Fc domain to become bivalent or multiple antibodies and thereby neutralize SARS-CoV-2^19,20^, we have successfully identified 7 monovalent Nbs with potent neutralizing activities. Nb11-59, at a concentration lower than 1 μg/mL, significantly inhibited the replication of authentic SARS-CoV-2 *in vitro*, and our monovalent Nb11-59 could be easily delivered by nebulization considering its small size and high stability. In addition, this molecule could be stably produced on a large scale at relatively low cost. Overall, these findings suggest that Nb11-59 may be a promising prophylactic and therapeutic during COVID-19 outbreaks.

## Results

### Identification of Nbs specific to SARS-CoV-2 spike RBD

In order to obtain Nbs with high affinity, specificity and good diversity, four camels were immunized with the recombinant RBD of SARS-CoV-2 spike. After seven injections at an interval of five days, the peripheral blood of each immunized camel was collected to construct the phage displayed library. The four libraries have more than 10^9 colony-forming units (CFU), and the insertion rate of each library was greater than 90%, indicating a high quality and good diversity of the four libraries. After three rounds of phage display biopanning, periplasmic extract ELISA (PE-ELISA) was performed to identify binding partners of RBD. The entire schedule was illustrated in Fig. 1.

**Fig. 1.**
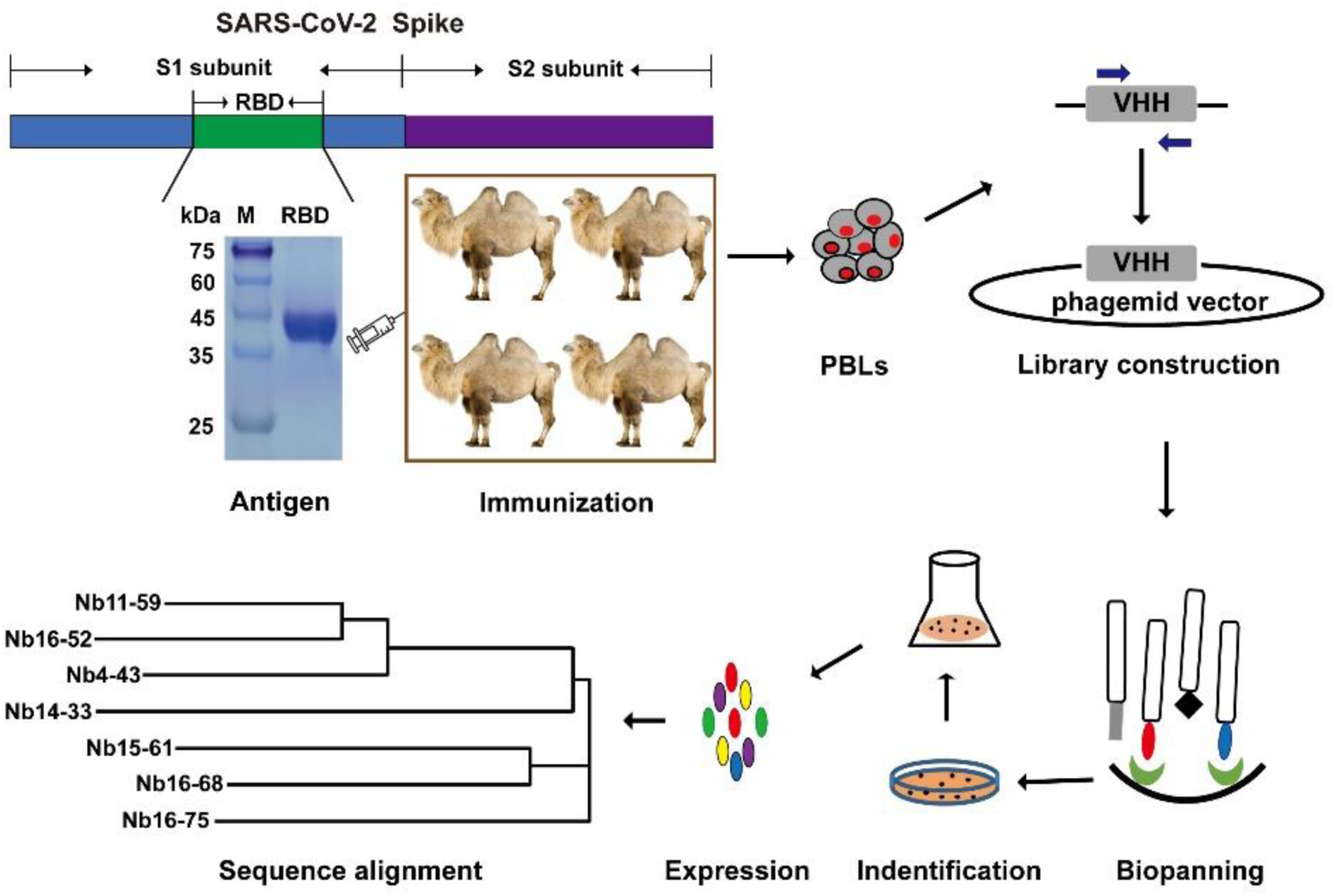
Schematic depicting the immunization and screening strategy that were used to isolate SARS-CoV-2 Spike RBD-directed Nbs.

In PE-ELISA assay, as shown in Fig. 2A, 690 of 1600 clones were identified as positive Nbs using the criterion of a binding ratio higher than 3 compared to negative control, and most of the ELISA-positive colonies showed high binding activity to the RBD (Fig. 2A). All the positive 690 Nb colonies were sequenced and the repeat sequences were removed based on the alignment of amino acid sequences. Of note, 381 RBD-specific clones with distinct sequences were identified and the phylogenetic tree analysis was performed based on the amino acid sequences of 381 Nbs by Clustal Omega (Fig. 2B). Sequence analysis of the 381 distinct Nbs demonstrated substantial sequence diversity in terms of both composition and length of these sequences, which offers us more options and potential candidates in the isolation of neutralizing Nbs.

Given the RNA virus nature of SARS-CoV-2, and the reported mutations of SARS-CoV-2-RBD in different countries, cross-activity of 381 Nbs against various RBD mutants was also studied through PE-ELISA. The detailed information was provided in Fig. S1. Of the 381 Nbs, 229 were identified to bind all the 8 mutants (V483A, V367F, N354D, V341I, Q321L, K378R, Y508H, H519P), and the remaining Nbs bound at least 1 mutant. These results demonstrated that these Nbs library could be a valuable screening resource for effective diagnostic and therapeutic reagents for COVID-19.

**Fig. 2.**
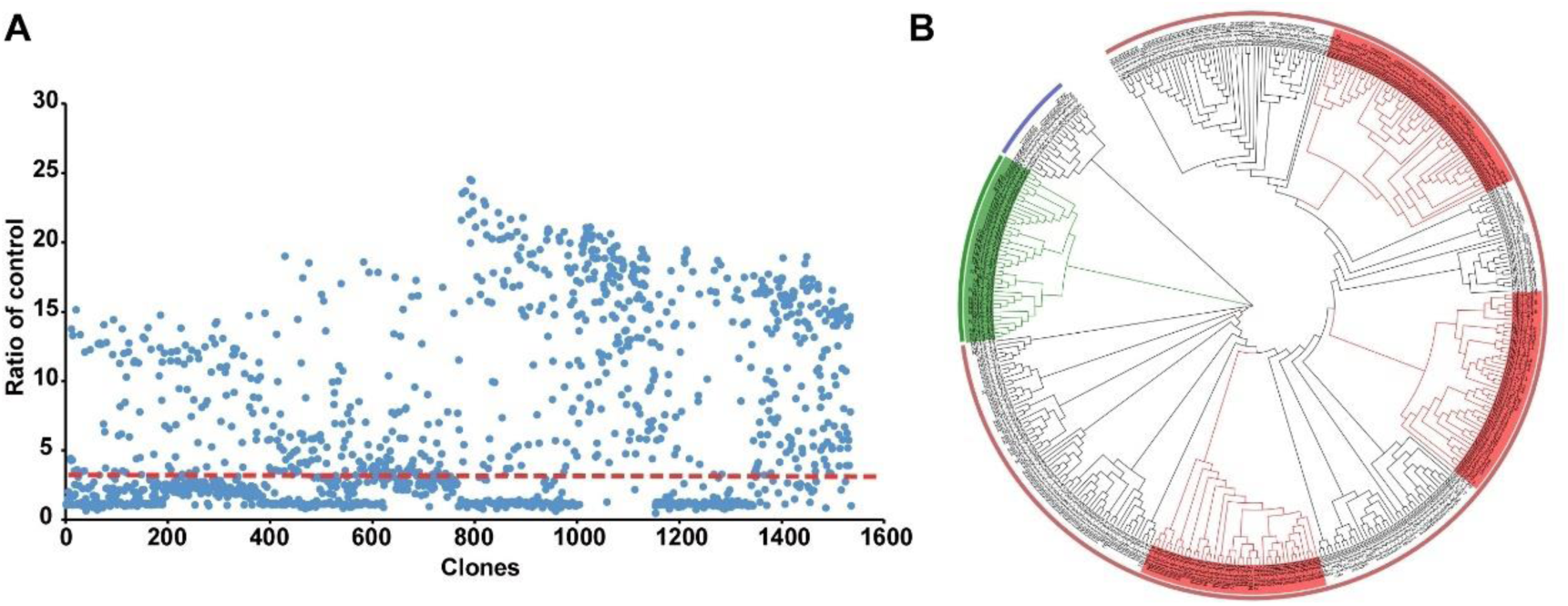
Identification of Nbs specific to SARS-CoV-2-RBD. (A) PE-ELISA was performed to identify positive clones. The clones with binding ratio higher than 3 was considered as positive clones. (B) Phylogenetic tree of the isolated SARS-CoV-2-RBD-directed Nbs, based on the neighbor joining method.

### Identification and characterization of Nbs with SARS-CoV2-RBD/ACE2 blocking activity

According to the cell-based blockade assay, 32 Nbs derived from lysis supernatants of 381 Nbs were considered as functional candidates, which have the ability to inhibit the interaction between RBD and ACE2, with the blocking rate higher than 15%. As indicated in Fig. 3A and B, Nb8-87, Nb13-58 and Nb11-59 exhibited blocking rates of 16.2%, 50.4% and 98.9%, respectively, which represented low, middle and high inhibitory functions. In order to further study the function and characterization of the 32 Nbs, we purified these Nbs through Ni-NTA affinity chromatography. SDS-PAGE analysis of the 32 Nbs were shown in Fig. 3C. The molecular weight (MW) of the 32 Nbs were all around 15 kDa, which is consistent with theoretical value. The Nbs are of high purity and could be used to study the blocking functions.

**Fig. 3.**
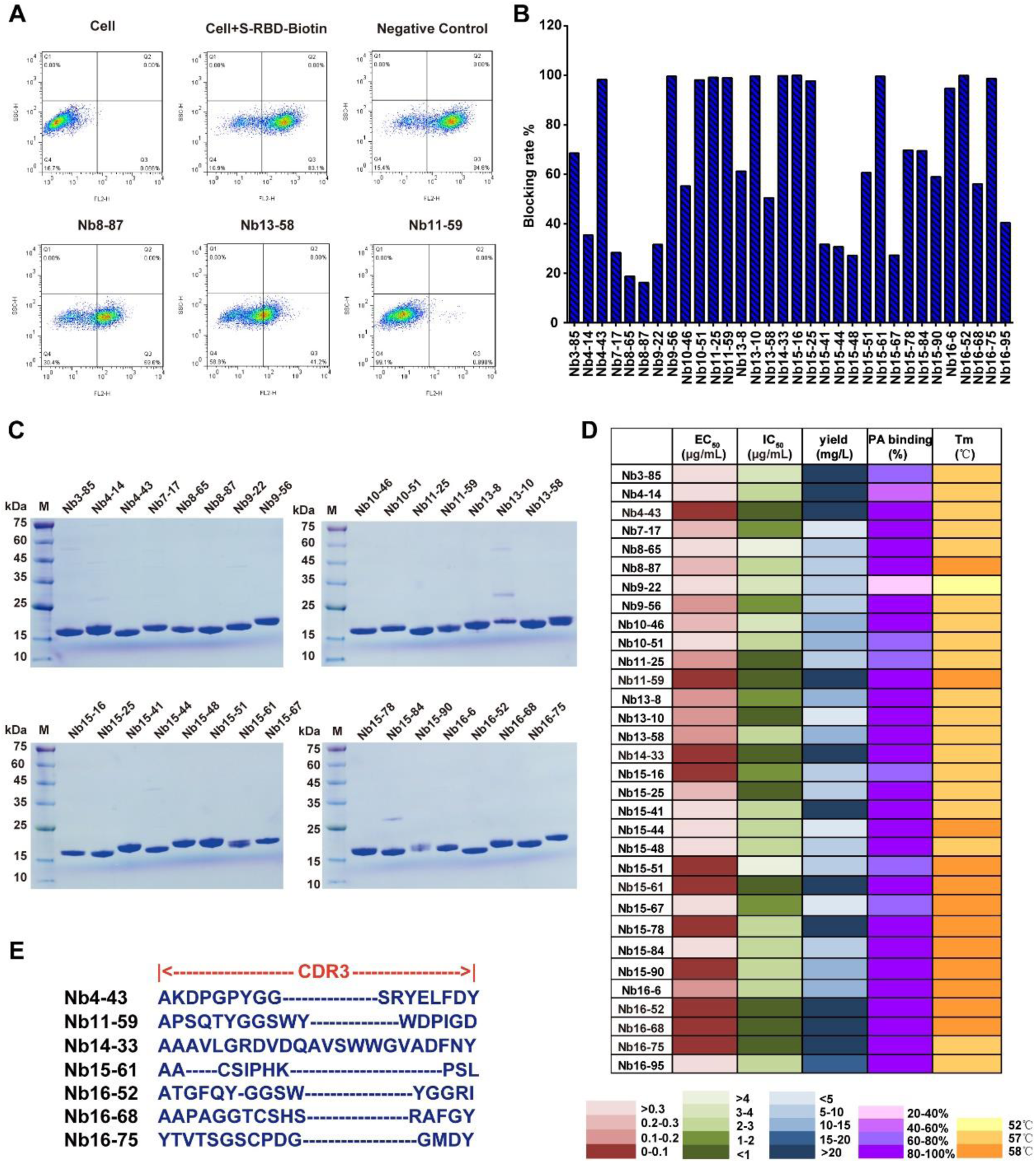
Identification and characterization of SARS-CoV-2-RBD/ACE2 blocking Nbs. (A) Examples are given to illustrate the results of screening by FACS. (B) Screening for SARS-CoV-2-RBD/ACE2 blocking Nbs. (C) Purification of blocking SARS-CoV-2-RBD Nbs and SDS-PAGE was used to check the purity of these Nbs. (D) Characterizations of blocking Nb candidates including EC_50_, IC_50_, yield, Protein A binding and Tm were displayed. (E) Sequences of CDR3 region of 7 selected SARS-CoV-2-RBD Nbs.

Next, we comprehensively studied the characterizations of Nbs including binding affinity, blocking activity, protein A binding activity and Tm value. The results showed that 17 of 32 Nbs bound to SARS-CoV-2-RBD with EC_50_ lower than 0.2 μg/mL, and 15 Nbs candidates exhibited obvious blockade effect towards SARS-CoV-2-RBD/ACE2 interaction, with IC_50_ lower than 2 μg/mL, among which the minimal IC_50_ was 0.580 μg/mL. Meanwhile, all Nb candidates could bind to Protein A agarose beads, with over 70% candidates showing binding efficiency higher than 80%, which allows for a relatively easy purification process in the following large-scale manufacturing. Additionally, the Tm values of these Nb candidates ranged from 52°C to 58°C (Fig. 4D, supplementary Fig. 2). With a comprehensive evaluation of various parameters, we eventually narrowed down to 7 Nbs candidates including Nb4-43, Nb11-59, Nb14-33, Nb15-61, Nb16-52, Nb16-68 and Nb16-75. These candidates have EC_50_ and IC_50_ lower than 0.2 μg/mL and 1μg/mL, respectively, as well as yield higher than 20 mg/L. Their PA binding efficacy is higher than 80% and their Tm is higher than 57°C. The sequences of these 7 candidates were analyzed, especially the amino acid sequences of complementarity determining region 3 (CDR3), which was considered as the key and most variable region of antibody. The results were demonstrated in Fig. 3E. The diversity in both composition and length among these 7 Nbs provides more possibilities and alternatives to identify neutralizing Nbs.

**Fig. 4.**
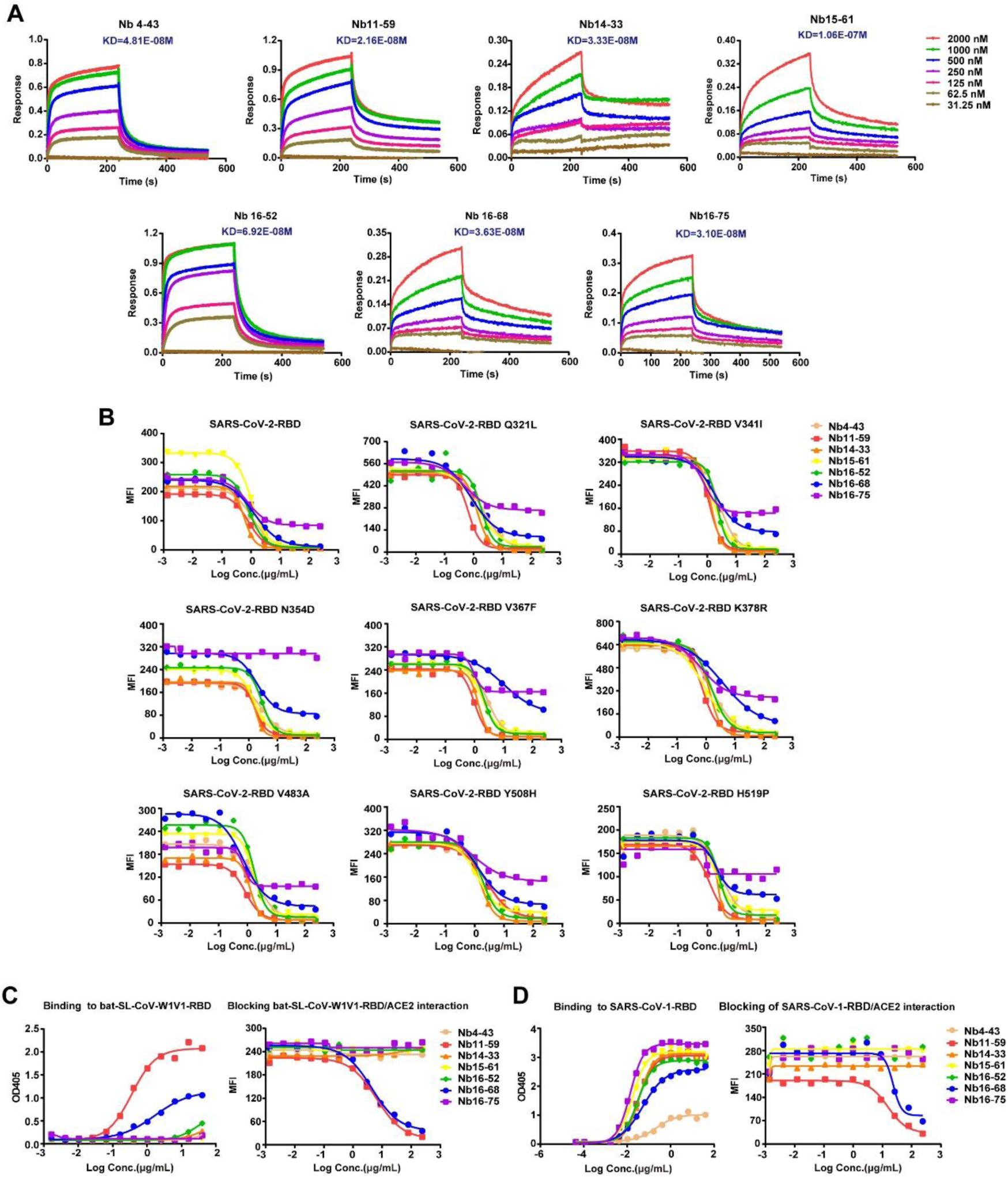
The affinity and blocking capacity of 7 Nbs towards SARS-CoV-2-RBD different mutants and other coronovirus species. (A) Affinity of selected candidates was measured by BLI. (B) The blocking activity against SARS-CoV2-RBD mutants/ACE2 interaction were determined by FACS. (C) The binding and blocking activities of selected 7 candidates towards bat-SL-CoV-WIV1-RBD were detected through ELISA and FACS, respectively. (D) The binding and blocking activities of selected 7 candidates towards SARS-CoV-1-RBD were detected through ELISA and FACS, respectively.

### Functional study of 7 Nbs against SARS-CoV-2-RBD including mutants

The seven Nbs which represented excellent binding capacity to RBD and outstanding ability to block infection were chosen to conduct comprehensive functional study on SARS-CoV-2-RBD or different RBD mutants. Firstly, the Biolayer Interferometry (BLI) based assay was carried out to measure the binding kinetics, and the results showed that all of these Nbs bound to SARS-CoV-2-RBD with high affinity. The Kd values ranged from 21.6 nM to 106 nM (Fig. 4A). Almost all of the 7 candidates could also bind to eight SARS-CoV-2 spike RBD mutants, including Q321L, V341I, N354D, V367F, K378R, V483A, Y508H and H519P variants, which are circulating in the US, England, France and China (date not shown). Furthermore, these 7 Nbs could block the interaction between ACE2 and eight different SARS-CoV2-RBD variants (Fig. 4B), with an exception of Nb16-75 which cannot bind N354D variant, suggesting N354 as an important recognizing epitope for Nb16-75 (Fig. 4B).

Given that the pandemic caused by coronavirus may re-arise frequently in the next few decades, it is necessary to find anti-coronovirus drugs with broad-spectrum neutralizing activity. Hence, these 7 candidates were analyzed for their neutralizing activity for other closely related betacoronaviruses species, bat-SL-CoV-WIV1-RBD and SARS-CoV-1-RBD. Of note, two Nbs, Nb11-59 and Nb16-68, demonstrated neutralizing potency against both bat-SL-CoV-WIV1-RBD and SARS-CoV-1-RBD. (Fig. 4C and Fig. 4D).

### Nb16-68 and Nb11-59 exhibit potent neutralizing activities against authentic SARS-CoV-2

Next, the neutralizing activity of 7 Nbs was determined against authentic SARS-CoV-2 *in vitro*. As indicated in Fig. 5A, all 7 Nbs exhibited a potent neutralizing ability against authentic SARS-CoV-2 in the PRNT at the concentration of 50 μg/mL and 5 μg/mL, with inhibitory rate of higher than 60% (Fig. 5A and 5B). Among these 7 Nbs, Nb16-68 and Nb11-59 showed the best neutralization activities. These two Nbs were serially diluted in 3-fold and their antiviral potencies were determined. Nb16-68 and Nb11-59 showed a potent antiviral activity against SARS-CoV-2 in Vero E6 cells in a dose-dependent manner, as judged by decrease of phage counts, with ND_50_ of 2.2 μg/mL and 0.55 μg/mL, respectively. (Fig. 5C and 5D). Overall, Nb16-68 and Nb11-59 might be novel potent candidates for COVID-19 treatment.

**Fig. 5.**
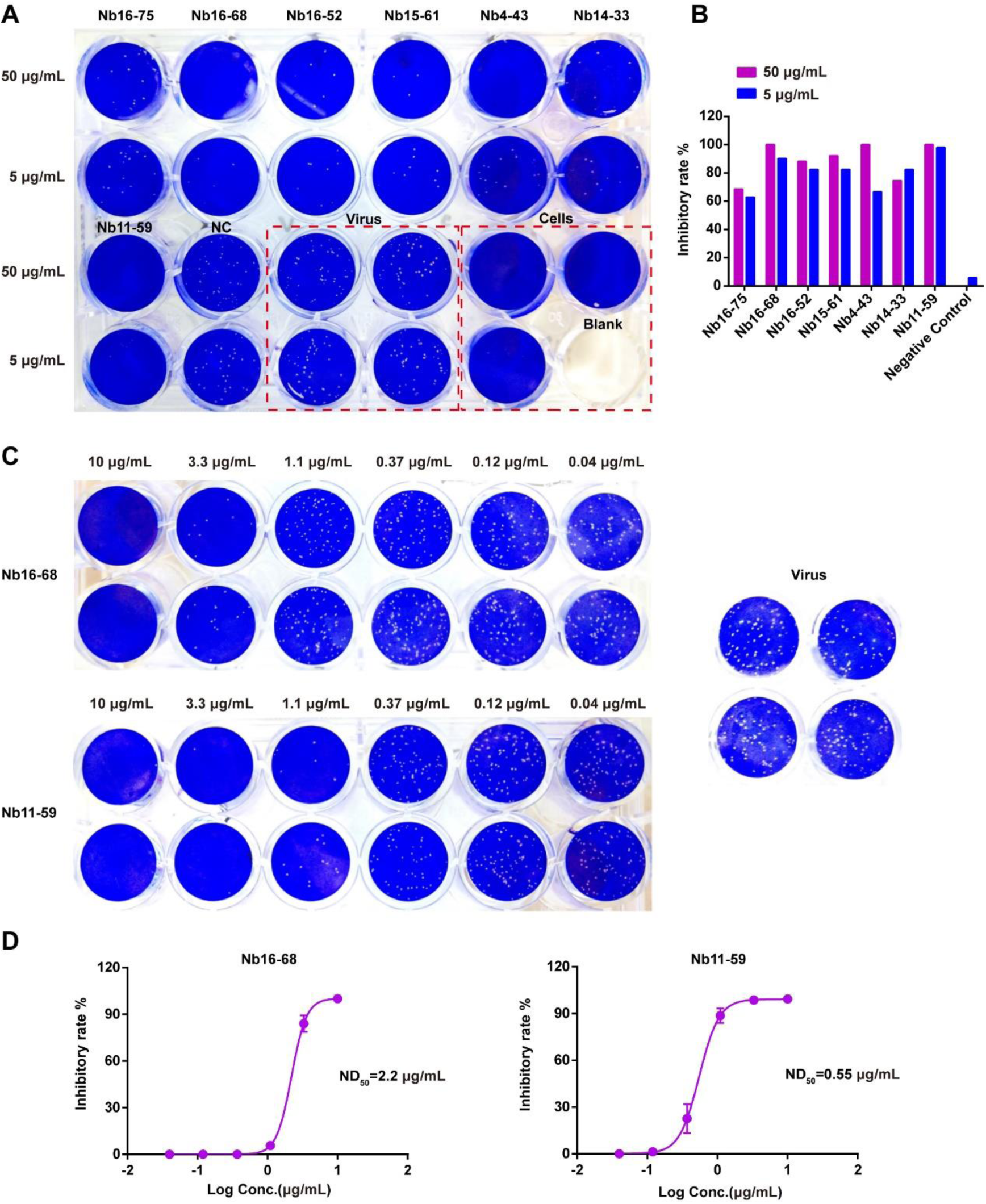
Evaluation of neutralizing potential of 7 Nbs using plaque reduction neutralization test. (A) Plaques formed in Vero E6 cells inoculated with 100 PFU SARS-CoV-2 and 10-fold diluted Nbs. Cry1B Nb was used as the negative control. (B) Inhibitory rate of 7 Nbs against authentic SARS-CoV-2 at the concentration of 50 μg/mL and 5 μg/mL. (C) Plaques formed in Vero E6 cells inoculated with 100 PFU SARS-CoV-2 plus 3-fold diluted Nb16-68 or Nb11-59 mixture. (D) 50% neutralizing dose (ND_50_) of Nb16-68 or Nb11-59 were determined. The results shown are mean value ± SD.

### Nb11-59 is predicted bound to the RBD region of the Spike protein

Considering Nb11-59 exhibited the best neutralization activity towards authentic SAR-CoV-2, we selected Nb11-59 for further structure prediction analysis. For Nb11-59, crystal structure of MT-SP1 in complex with Fab Inhibitor E2 (PDB 3BN9)^21^ with sequence identity of 82.26% was used as the template. The homology modeling obtained GMQE values^22^ (close to 1 means very close to the experimental results) of 0.97 for Nb11-59. QMEAN values (over than -4 is acceptable) is 0.55 for Nb11-59 (data not shown). Results indicate that the homology modeling of Nb11-59 is favorable and can be used for further prediction of the binding interaction with SARS-CoV-2 Spike protein.

Next, according to the results of Nb conformation and its binding epitope prediction, Nb11-59 binds to the RBD region of the Spike protein. At the binding interface of complex, Tyr449 as well as amino acids ranging from 489 to 501, which are within the Spike protein RBD region, may contribute to the binding and are regarded as critical residues for the binding affinity. The predicted residues may contribute to the binding interaction of the Nb and Spike protein, and can be confirmed by further mutagenesis analysis.

### Large-scale production, stability analysis and nebulization of humanized Nb11-59

Nb11-59 was humanized according to our previous studies^17,23^. For future clinical application, large-scale production of SARS-CoV-2 Nb with low cost is of great necessity. Therefore, we tried to express the humanize Nb11-59 (HuNb11-59) in *Pichia pastoris* by fermentation. After initial cloning into *P. pastoris*, about 100 transformants of HuNb11-59 were tested for small-scale expression test, and the colonies with highest expression were chosen for further fermentation. During the fermentation process in 7L fermenter, the yield of HuNb11-59 protein increased. Surprisingly, after 213 hours of induction, the yield of target protein in fermentation supernatant has even reached 20 g/L (Fig. 7A), which is, to our knowledge, the highest expression of Nbs among the yield of Nbs that have been reported. After affinity chromatography and hydrophobic chromatography purification procedures, HuNb11-59 exhibited high purity through SDS-PAGE analysis (Fig. 7B), and the specific purity of Nb was 99.36%, 99.55% and 95.95%, respectively, through SEC-HPLC, AEX-HPLC and RP-HPLC analysis (Fig. 7C), which means the commercial manufacturing and purification process have been successfully developed and validated, and the key methods in analytical quality characterization have been already established.

**Fig. 6.**
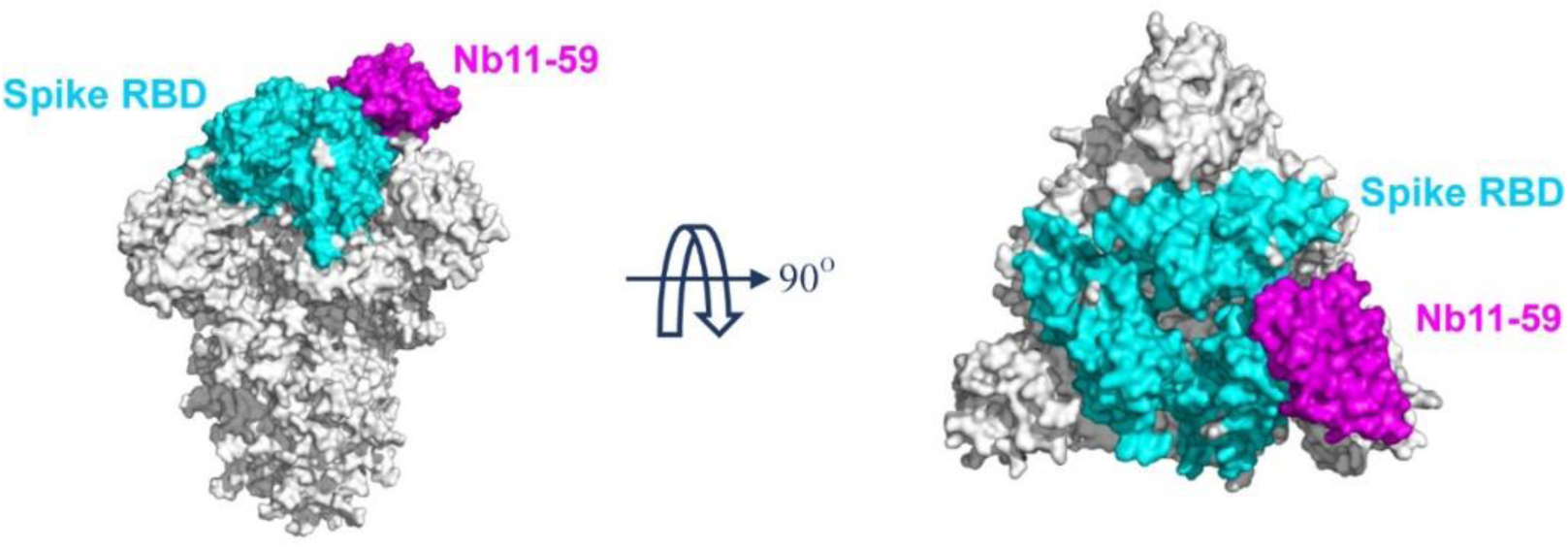
The binding model of Nb11-59 to Spike protein. Nbs and Spike RBD were represented as purple and blue surface, respectively.

**Fig. 7.**
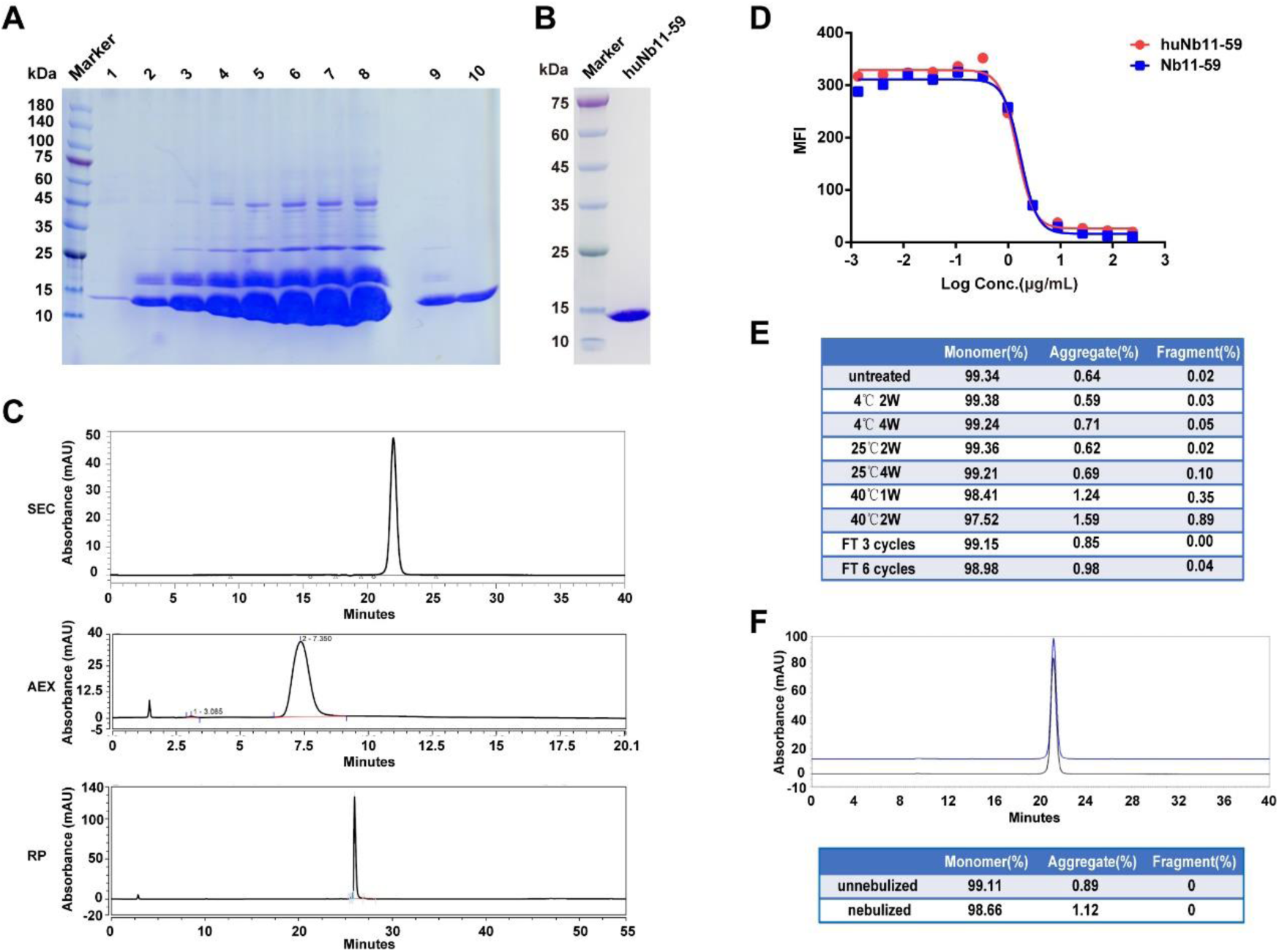
The large-scale production and preliminary druggability analysis. (A) The yield of HuNb11-59 in fermentation tank was determined at indicated times, which were detected by SDS-PAGE assay, with 1g/L Nb11-59 as the standard protein. lanes 1-8, 10 µL supernatants after 0, 24, 48, 72, 120, 159.5, 183.5 and 213 h of induction; lane 9, 10 µL 1/20 diluted culture supernatant induced for 213 h; lanes 10, 10 µL 1g/L high purity target protein. (B) The purity of HuNb11-59 was determined through SDS-PAGE, following affinity chromatography and hydrophobic chromatography. (C) The purity of HuNb11-59 was determined through SEC-HPLC, AEX-HPLC and RP-HPLC analysis. (D) The blocking activity of Nb11-59 and HuNb11-59 against SARS-CoV-2-RBD/ACE interaction was determined by FACS. (E) The stability of HuNb11-59 under different temperatures and of freeze-and-thaw cycles was detected by SEC-HPLC. (F) The purity of nebulized HuNb11-59 formulation was detected by SEC-HPLC.

Following the successful purification of HuNb11-59 from yeast fermentation, FACS based SARS-CoV-2-RBD/ACE2 blocking activity assay was conducted for both Nb11-59 and HuNb11-59. As shown in Fig. 7D, Nb11-59 and HuNb11-59 demonstrated similar profile in SARS-CoV-2-RBD/ACE2 neutralization. Furthermore, the stability of HuNb11-59 was detected at various conditions. The percentage of monomer was rather stable, maintaining above 99% when stored at 4°C or 25°C for up to 4 weeks. While being left at 40°C for 2 weeks, only 0.89% degradation and 1.59% aggregation was found (Fig. 7E). At the same time, a total of 6 cycles of freeze-and-thaw showed no effect on the percentage of monomer (Fig. 7E). These stability data showed great drug stability of HuNb11-59, which ensured the quality, safety and efficacy of the drug product. In addition, in order to make HuNb11-59 administered as an inhaler directly to the lung, great efforts were made to find out a proper formulation for the nebulization of HuNb11-59. The SEC-HPLC results indicated that the nebulization under our proposed formulation conditions did not siginificantly undermine the stability of HuNb11-59, with only a very minor proportion of aggregates found. (Fig. 7F). The results demonstrated that we have creatively developed the first Nb drug against SARS-COV with the delivery manner of respiratory drug. The novel way of drug delivery may offer patients more convenient drug administration and better drug absorption effects.

## Discussion

In this study, we reported Nb phage display libraries derived from four camels immunized with the SARS-CoV-2 spike RBD, from which 381 Nbs were identified to recognize the RBD including the eight variants of SARS-CoV-2, suggesting that the these RBD specific Nbs collection could be very valuable in the development of diagnostic and therapeutic reagents for COVID-19 infection. Currently, Nb based diagnostic methods have not been reported yet. It would be very meaningful to develop SARS-CoV-2 virus fast-tracking techniques with Nbs^24,25^. In addition to the diagnostic value, Nbs could also be developed as PET-CT probes for their good tissue infiltration ability and short half-life in vivo^26,27^. The large RBD specific Nb library in this study would facilitate the screening for COVID-19 preventive and therapeutic candidates.

We identified a potently neutralizing molecule of Nb11-59, which recognizes the wildtype and eight kinds of mutants of RBD proteins. Nb11-59 was capable of inhibiting the replication of authentic SARS-CoV-2 *in vitro* with ND_50_ of 0.55 μg/mL, which is similar to or lower than the ND_50_ or IC_50_ values of several reported mAbs isolated from human B cells^4,28,29^. Importantly, Nb11-59 herein demonstrates neutralizing abilities more than 4 fold higher than other reported Nbs, including n3130 and n3088 with IC_50_ of 4.0 μg/mL and 2.6 μg/mL, respectively, against authentic SARS-CoV-2^30^. The published neutralizing Nbs towards SARS-CoV-2 were evaluated mainly in the pseudoviruse system, with IC_50_ ranging from 0.003 μg/mL to 12.32 μg/mL^19,31-34^. Of note, IC_50_ of molecules tested in the pseudoviruse system might be 2-100 fold lower than that in the authentic SARS-CoV-2^29,32^. Thus, Nb11-59 might have higher neutralizing abilities against authentic SARS-CoV-2 compared with most of other reported Nbs without Fc fusion.

With the continuing COVID-19 epidemic, the resistance to any potential antiviral therapeutic would be the focus due to the rapid mutation of viral pathogens, especially under the selective pressure of antiviral drugs. Indeed, naturally occurring RBD variations, such as V483A, V367F and V341I variant, have been circulating in some regions of the world^35^. A number of drugs, especially RBD-directed antibodies with potent neutralizing activities against SARS-CoV-2 have been described^35^. Nevertheless, there are few reports about antibodies that can neutralize a wide range of SARS-CoV-2 variants. Herein, we showed that 7 RBD-directed Nbs identified were capable of blocking the interaction between ACE2 and the eight RBD-mutant variants, with Nb11-59 showing the most potent blocking activities. In addition, these molecules also demonstrate blocking activities against SARS-CoV-1 and bat-SL-CoV-WIV1. Further study needs to be performed for evaluating the neutralizing activities against authentic SAR-CoV-2 variants. Notably, the variants resistant to these molecules would likely arise under the selective pressure although they might have broad spectrum activities against the different SAR-CoV-2 variants. Non-competing antibody cocktail treatment might be a good strategy to prevent rapid mutational escape of SARS-CoV-2^36^.

The development of vaccines and mAbs represents a promising strategy for combating COVID-19 infections; however, they might induce antibody-dependent enhancement (ADE), an undesirable phenomenon that leads to increased infectivity and virulence, which has been observed in the infection of SARS-CoV-1, MERS COV, HIV and Ebola viruses with the treatment of non-neutralizing or neutralizing mAbs^37-40^. It is known that ADE is likely to be associated with Fc domain of antibodies. Despite the substantial attempts made at the modifications of Fc domains to avoid the emergence of ADE, the expected results have not been achieved^41,42^. Because of lacking Fc domain, Nbs or VHHs represent ideal neutralizing antibodies against viral infections. A typical example is Nb ALX-0171 with potent neutralizing activities against respiratory syncytial virus (RSV), which is currently in phase II clinical trials^43^. Therefore, the development of neutralizing Nbs that do not contain Fc domains has significant advantages in the fight against COVID-19.

Considering their small size and biophysical properties, single domain antibodies have multiple advantages in the treatment of respiratory tract infectious diseases. First, single domain antibodies could be produced on a large scale at low cost. We showed that Nb11-59 expression could reach 20 g/L through fermentation using the yeast expression system, which means that it could be rapidly and widely used as a preventive or therapeutic molecule. Second, Nb11-59, as a single domain antibody with small size, could be delivered to the site of infection via inhalation, which is supported by its high stability at the temperature ranging from 4°C to 40°C, and its consistent post-nebulization stability profile. Because of the highly infectious and continuous outbreak of COVID-19, the inhaled delivery of Nb11-59 is likely effective in controlling the spread of the virus. Taken together, Nb11-59 is a very potential therapeutic molecule against COVID-19, and it is worthy of further research to facilitate rapid clinical development.

## Methods

### Cells and Viruses

The HEK 293 and Vero E6 cells were obtained from the American Type Culture Collection (ATCC) and China Center for Type Culture Collection (CCTCC), respectively. The Vero E6 cells were grown in Dulbecco’s Modified Eagle Medium (DMEM) (Gibco, GrandIsland, NY, USA) supplementing with 1% Penicillin-Streptomycin (10,000 U/mL) (Gibco) and 10% FBS (Gibco). The HEK 293 were grown in DMEM supplemented with 1% Penicillin-Streptomycin and 10% FBS for Nbs functional analysis, and in CD05 (OPEM, SH, CN) for proteins production. The SARS-CoV-2 strain IVCAS 6.7512 was offered by the National Virus Resource, Wuhan Institute of Virology, Chinese Academy of Science^44^. All processes in this study involving authentic SARS-CoV-2 were performed in a biosafety level 3 (BSL-3) facility. The illustrations of authentic SARS-CoV-2 in Graph Abstract was derived from the source page: *Desiree Ho for <u>the Innovative Genomics Institute</u>.*

### Generation of SARS-CoV-2 spike RBD wildtype and mutant proteins

The coding sequence of SARS-CoV-2 spike RBD was achieved from the UniProt website (https://www.uniprot.org/). The RBD wildtype and mutant proteins of SARS-CoV-2 spike with 10×his tag at N terminal were expressed in HEK 293 and purified with Ni-NTA affinity columns (Qiagen, Hilden, Germany).

### Camel immunization and phage display library construction

The RBDs of SARS-CoV-2 spikes were used as antigens to conduct camel immunization, then the immunized phage display library was generated according to our established methods^45,46^. Briefly, four camels were injected with RBD antigens once every 5 days for a total of seven times. The peripheral blood from these immunized camels was collected for lymphocytes cells extraction and phage libraries were constructed. All procedures were performed according to the Health guide for the care and use of laboratory animals.

### Biopanning of Nbs against SARS-CoV-2 spike RBD

Phage display technology was used to perform Nbs biopanning. RBD with his tag was diluted in 100 mM NaHCO_3_ and coated in microplate overnight. After blocking with BSA, the phages from the immunized library were added into wells for incubation. The specific phages were eluted after strict and harsh wash procedure to remove the unspecific binders. The specific phages would be enriched and prepared for the next round of biopanning.

### Periplasmic extract ELISA (PE-ELISA)

In order to identify positive clones, 400 clones from each of the 4 libraries were randomly picked and expressed in microplate for PE-ELISA verification. Each clone was cultured in Terrific Broth medium (Invitrogen) for 3 h and induced by 1 mM IPTG (Sigma-Aldrich, USA) overnight. After an osmotic shock, the supernatants were transferred into wells of the microtiter plate coated with SARS-CoV-2-RBD-His in advance. The mouse anti-HA antibody (Covance, Princeton, NJ, USA) and goat anti-mouse IgG-alkaline phosphatase (Sigma-Aldrich) were added to the wells for incubation successively. Then, 405 nm absorbance was read by the microplate reader (Bio-Rad, Hercules, CA, USA) after the substrate of alkaline phosphatase was added. The positive clones were identified when the ratios were higher than 3.

### Nbs expression and purification

After sequencing the selected clones, the SARS-CoV-2-RBD specific clones were amplified, then the plasmids of these candidates were extracted and transformed into *Escherichia coli* strain WK6, Ni-NTA affinity columns (Qiagen, Hilden, Germany) were used in purification of Nbs. Purified Nbs were confirmed through sodium dodecyl sulfate polyacrylamide gel electrophoresis (SDS-PAGE).

### Phylogenetic tree

The multiple alignment and the construction of the tree was made by Clustal Omega, and pair-wise distance was calculated by the strategy of Neighbor-joining^47^.

### Affinity determination

The kinetics of Nbs binding to SARS-CoV-2-RBD antigen were performed by biofilm interferometry (BLI) with a Fortebio’s Octet RED96 instrument (ForteBio, Menlo Park, CA, USA). Briefly, the diluted Nbs (10 μg/mL) were coupled to protein A biosensors and then incubated with a series diluted SARS-CoV-2-RBD, followed by dissociation in PBST. The binding curves were fit in 1:1 binding model by Octet Data Analysis software 9.0. The association and dissociation rates were monitored and the equilibrium dissociation constant (KD) was determined.

### Activity assay of Nbs binding to SARS-CoV-2-RBD

To determine the activity of anti-SARS-CoV-2-RBD Nbs binding to the antigen of RBD, serial dilution of anti-SARS-CoV-2-RBD Nbs were incubated with 1 μg/mL SARS-CoV-2-RBD-His coated on the 96 microtiter plate wells for 1 h. Next, the plates were incubated with the mouse anti-HA antibody followed by goat anti-mouse IgG-alkaline phosphatase (Sigma-Aldrich). The absorbance at 405 nm was read by the microplate reader (Bio-Rad, Hercules, CA, USA), and the 50% effective concentration (EC_50_) was determined.

### Activity assay of Nbs blocking SARS-CoV-2-RBD/ACE2

To determine the activity of anti-SARS-CoV-2-RBD Nbs blocking SARS-CoV-2-RBD/ACE2 interaction, ACE2/HEK 293 stable cell line was constructed using a lentiviral packaging system. 3×10^5^ ACE2/HEK 293 cells were incubated with the 2.5 μg/mL purified SARS-CoV-2-RBD labeled with biotin, and a gradient concentration of anti-SARS-CoV-2-RBD Nbs, followed by staining with streptavidin-PE (eBioscience, San Diego, CA, USA). The signals were measured by BD FACS Calibur instrument (BD Biosciences, Franklin Lakes, New Jersey, USA), and the 50% inhibitory concentration (IC_50_) was determined.

### Protein A binding assay

1 mg of Nbs were incubated with 20 μL of Protein A chromatography resin at room temperature for 30 min. Then the mixtures were centrifuged and the protein concentration of supernatants were determined by NanoDrop. The protein A binding percentage was calculated.

### Tm measured by thermal shift assay (TSA)

To determine the thermal stability of Nbs. TSA were performed in a high-throughput manner (96-well plate) with a Q-PCR device, and 0.1 mg/mL of Nbs were mixed with 10×Sypro Orange protein Gel stain in each tube. The program consisted of the following steps: heat to 25°C at a ramp rate of 4.4°C/s and hold for 25 s; heat to 98°C at a continuous ramp rate of 0.1°C/s; then cool to 25°C and hold for 10 s.

### Authentic SARS-CoV-2 plaque reduction neutralization test (PRNT)

PRNT was used to test the blockade of SARS-CoV-2 virus attachment by Nbs. Briefly, Vero E6 cells were seeded into 24-well culture plates at 1.5×10^5^ per well and incubated at 37°C in 5% CO_2_ overnight. Nbs were serially diluted in DMEM supplemented with 2% FBS. The diluted SARS-CoV-2 suspension containing 300 plaque-forming units (PFU)/mL was added into Nbs, and then incubated at 37°C for 1 h. 200 μL of the Nbs-virus mixture was added into a 24-well culture plate containing Vero E6 cells. In addition, cells infected with 150 PFU/mL of SARS-CoV-2 and those without the virus were applied as positive and negative controls, respectively. After incubation for 1 h at 37°C, the Nbs-virus mixture was removed from Vero E6 cells followed by 500 μL of DMEM with 2% FBS and 0.9% carboxymethyl cellulose (Promega) was overlaid. For further incubation at 37°C in 5% CO_2_ for 3 days, plaques were stained by 0.5% crystal violet. Individual plaques were counted for 50% neutralizing dose (ND_50_) calculation.

### Prediction of Nb Conformation and its binding interaction with SARS-CoV-2 Spike protein

The homology model of Nbs was constructed on the SWISS-MODEL online server (https://swissmodel.expasy.org/) based on the searched templates. The complex model of SARS-CoV-2 spike glycoprotein (PDB 6VXX)^48^ and Nb was built using Z-DOCK (http://zdock.umassmed.edu)^49^. Complex with the highest docking score was selected. The binding interaction model of Nb with SARS-CoV-2 Spike protein were generated by Pymol (https://pymol.org/2/).

### SEC-HPLC, AEX-HPLC and RP-HPLC assay

Nbs were capture and purified by protein A affinity chromatography, and then further purified by hydrophobic chromatography on AKTA pure 150 (GE Healthcare, Madison, WI, USA). The purity of Nbs was detected by size exclusion chromatography-high performance liquid chromatography (SEC-HPLC), anion exchange chromatography-high performance liquid chromatography (AEX-HPLC) and reversed phase chromatography-high performance liquid chromatography (RP-HPLC). SEC-HPLC analysis using Waters Acquity Arc system with AdvanceBio SEC 130A Columns (Agilent Technologies, Palo Alto, Calif, USA), AEX-HPLC analysis using Thermo Fisher Ultimate 3000 system with ProPacTM WAX-10 Columns (Thermo Scientific, Rockford, IL, USA) and RP-HPLC using Thermo Fisher Ultimate 3000 system with AdvanceBio RP-mAb Diphenyl Columns (Agilent Technologies).

### Large-scale expression in 7 L bioreactor

The vector containing VHH gene was linearized and transformed into *Pichia pastoris* to establish a high-efficiency expression system. Dozens of clones were screened in shake flasks. After identifying the high expression clone, large-scale expression was carried out in 7 L bioreactor according to the existing fermentation method of our company. During the fermentation process, fermentation broth was sampled at different fermentation time and centrifuged by 12000 rpm for 5 minutes. The supernatants of different fermentation time were tested by SDS-PAGE.

### Stability analysis

Formulated Nbs were stored at 4°C, 25°C and 40°C separately. The samples stored at 4°C and 25°C were examined for their purity at 2-week and 4-week time point by SEC-HPLC. The samples stored at 40°C were checked at 1-week and 2-week time point. The samples undergone 3 or 6 freeze-and-thaw cycles were also measured.

### Purity analysis after nebulization

Nbs purified from yeast fermentation at concentration of 10 mg/mL in a proper formulation were nebulized with a PARI eFlow rapid Nebulizer System (PARI GmbH, Starnberg, German). 8 mL of formulated Nbs were filled in the reservoir and operated for about 20 mins, the nebulized Nbs were collected with a 50 mL Centrifugal tube. The purity of the collected Nbs was performed on SEC-HPLC.

### Statistical analysis

Statistical analysis was performed using GraphPad Prism 6 software. Data are expressed as mean ± SD.

## Acknowledgements

We thank Jia Wu, Jun Liu and Hao Tang from BSL-3 Facility of Wuhan Institute of Virology for their essential support. We thank Center for Biosafety Mega-Science, Chinese Academy of Sciences and the National Virus Resource Center for resource support. We also thank Hongde Liu from Southeast University offers the assistance.

This work was funded by Shanghai Sailing Program (20YF1434300), National Natural Science Foundation of China (81902052) and the Natural Science Foundation of Hubei Province of China (2019CFA076).

## Author Contributions

Y.W. conceived and designed the experiments. J.G., L.M., G.L., M.Z., P.Q. and X.L. (Xiaofei Li) conducted the experiments, analyzed the data and prepared the manuscript, R.G. and H.Z. performed the neutralizing activity assay against authentic SARS-CoV-2, Y.Z. and Y.C. performed Nb conformation and its binding epitope prediction. W.J., H.Z., H.C. and X.L. (Xionghui Li) participated in library screening and functional study.

## Competing Interests

All commercial rights from this paper belong to Shanghai Novamab Biopharmaceuticals Co., Ltd.

**Correspondence** and requests for materials should be addressed to Y.W.

