## Supplementary Figure 1 for "A potent neutralizing nanobody against SARS-CoV-2 with inhaled delivery potential"

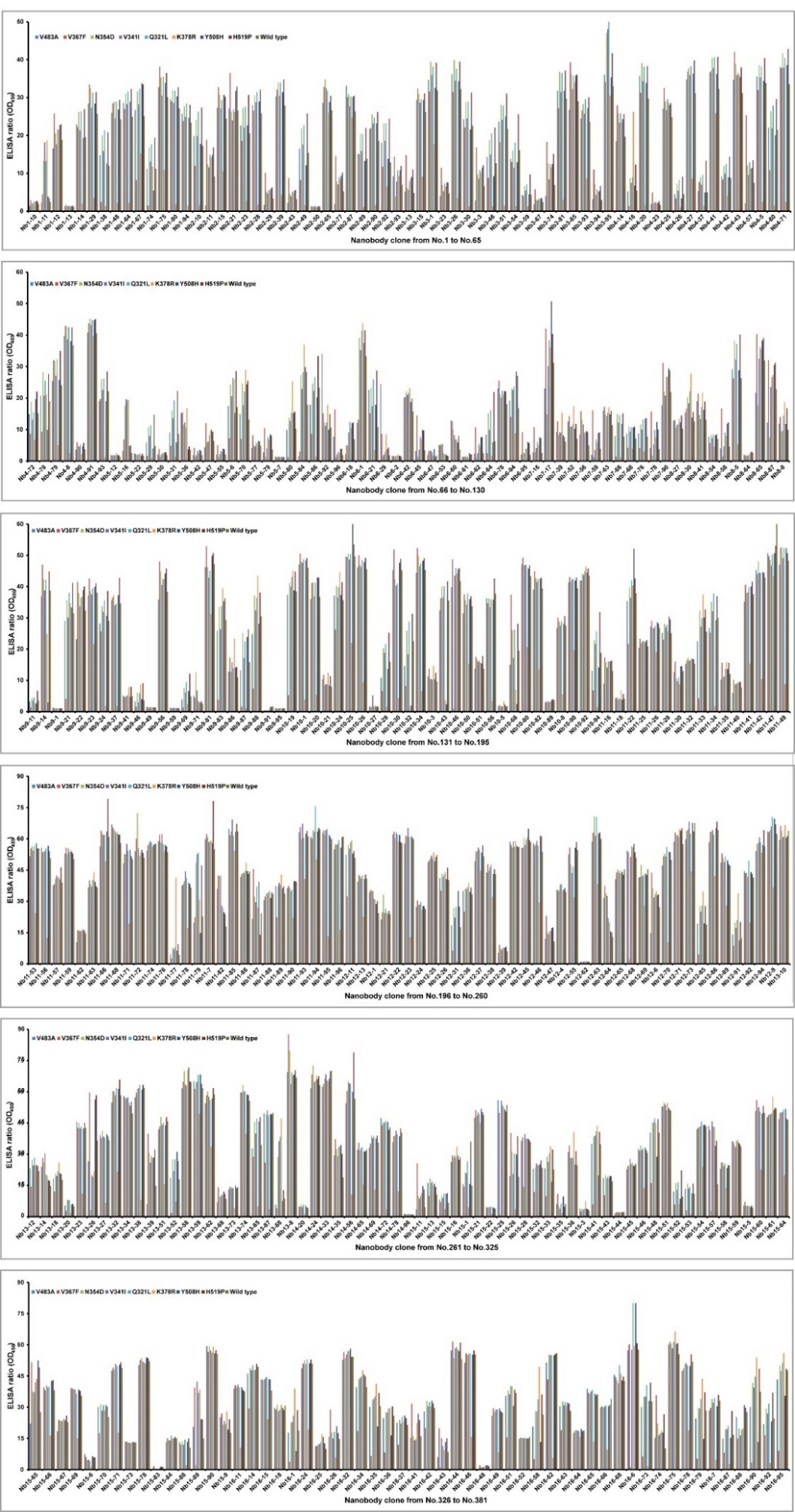

Supplementary Figure 1. The binding activity of positive colonies to different SARS-CoV-2-RBD mutants.
