## Supplementary figures and images for "A potent neutralizing nanobody against SARS-CoV-2 with inhaled delivery potential"

### Supplementary Figure 2

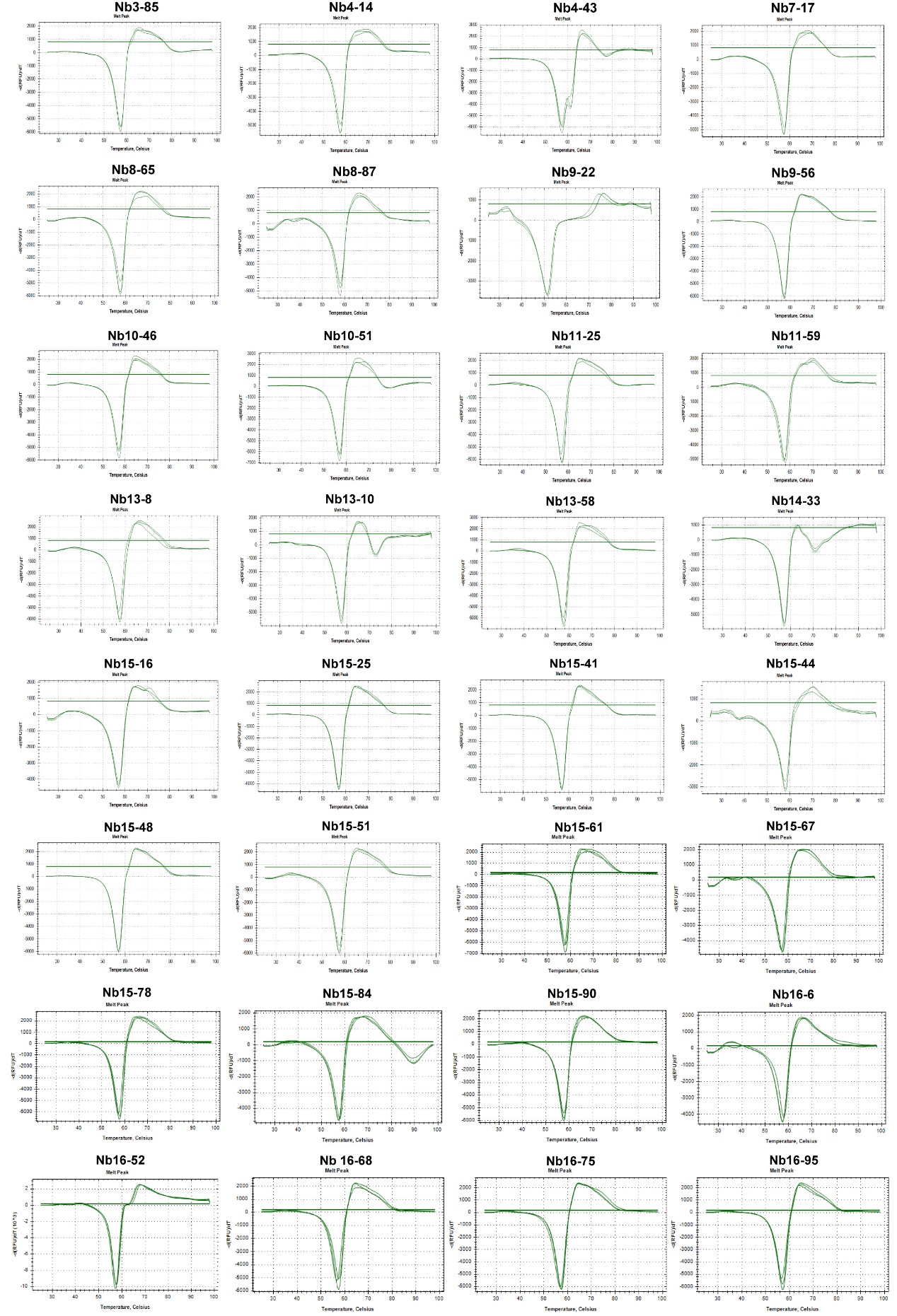

Supplementary Figure 2. The Tm value detection of the 32 purified SARS-CoV-2-RBD specific Nbs.
